# Metabolic depression in sea urchin barrens associated with food deprivation

**DOI:** 10.1101/2020.11.28.402156

**Authors:** Nathan B. Spindel, Lynn C. Lee, Daniel K. Okamoto

## Abstract

The proliferation of sea urchins can decimate macroalgal forests in coastal ecosystems, leading to persistent barren seascapes. While kelp forests are among the most productive ecosystems on the planet, productivity in these urchin barrens is dramatically reduced. Moreover, urchins inhabiting these food-depauperate barrens face starvation and many survive in these barrens for years or decades. Urchins in barrens can persist by eating food subsidies from drift algae, pelagic salps, tubeworms, as well as encrusting and filamentous algae, microbial mats, and slow-growing species resistant to herbivory. Despite both food from endogenous production and exogenous subsidies, many urchins in barrens likely experience prolonged food deprivation. This resource limitation may create a trade-off between reproduction and survival; for example, fecundity of purple sea urchins (*Strongylocentrotus purpuratus*) is 99.9% lower in barrens. Despite food constraints, red sea urchins (*Mesocentrotus franciscanus*), the dominant urchin species at our study sites, can live in excess of 100 years and barrens in Haida Gwaii, British Columbia (BC), Canada, have persisted for at least 143 years. While these phenomena are widespread and well documented, the bioenergetic adaptations that allow urchins to persist in these food-depauperate barrens remain poorly understood. To quantify habitat-specific differences in metabolic rates and energy reserves (as measured by gonadal mass), we conducted respirometry on and measured gonadal mass in *M. franciscanus* at three locations in BC inside and outside of adjacent kelp forest and barrens habitat. Here we demonstrate that *M. franciscanus* in barrens versus kelp forests have substantially lower energy reserves and, importantly, also exhibit dramatic reductions in size-specific resting metabolic rates (RMR), even after standardizing by metabolically active body mass. On average, gonadal mass was 44.6% lower and RMR scaled to metabolically active body mass was 40% lower in barrens urchins than in kelp forest urchins. Such a shift in metabolic rate may provide a mechanism that facilitates barren state stability over long time scales as *M. franciscanus* can lower energetic demands while they wait for small pulses of food, scrape by on low-productivity resources, and suppress recruitment of macroalgae for months, years, or decades.

---

The proliferation of sea urchins can decimate macroalgal forests in coastal ecosystems, leading to persistent barren seascapes. While kelp forests are among the most productive ecosystems on the planet, productivity in these urchin barrens is dramatically reduced (Filbee-Dexter and Scheibling 2014). Moreover, urchins inhabiting these food-depauperate barrens face starvation and many survive in these barrens for years or decades. Urchins in barrens can persist by eating food subsidies from drift algae (Rodríguez 2003, Vanderklift and Wernberg 2008, Britton-Simmons et al. 2009, Renaud et al. 2015, Quintanilla-Ahumada et al. 2018), pelagic salps (Duggins 1981), tubeworms (Spindel and Okamoto *personal observation),* as well as encrusting and filamentous algae, microbial mats, and slow-growing species resistant to herbivory (Ling and Johnson 2009, Filbee-Dexter and Scheibling 2014, Rasher et al. 2020). Despite both food from endogenous production and exogenous subsidies, many urchins in barrens likely experience prolonged food deprivation. This resource limitation may create a trade-off between reproduction and survival (Stearns 2000); for example, fecundity of purple sea urchins *(Strongylocentrotus purpuratus)* is 99.9% lower in barrens (Okamoto 2014). Despite food constraints, red sea urchins *(Mesocentrotus franciscanus),* the dominant urchin species at our study sites, can live in excess of 100 years (Ebert 2008) and barrens in Haida Gwaii, British Columbia (BC), Canada, have persisted for at least 143 years (Dawson 1880). While these phenomena are widespread and well documented, the bioenergetic adaptations that allow urchins to persist in these food-depauperate barrens remain poorly understood. Here we demonstrate that *M. franciscanus* in barrens versus kelp forests have substantially lower energy reserves (as measured by gonadal mass) and, importantly, also exhibit dramatic reductions in size-specific resting metabolic rates (RMR), even after standardizing by metabolically active body mass.

Animals often cope with severe food deficiencies by modifying their locomotive activity, utilizing reversible energy reserves, and/or increasing metabolic efficiency (McCue 2010). For example, green urchins *(Strongylocentrotus droebachiensis)* may invest energy in active searching for the potential return of richer pastures (Scheibling et al. 2020), while *M. franciscanus* may maintain a more sedentary lifestyle to conserve energy until food subsidies become available (Lowe et al. 2015). Urchins may resorb energy reserves stored in gonad tissues (Carey et al. 2016) and other tissues may lose mass because rates of cell loss exceed rates of proliferation (Secor and Carey 2011); such reductions in biomass can reduce energetic maintenance costs. Finally, starving animals can also reduce metabolic costs by downregulating cellular-level demand for and supply of ATP (Staples and Buck 2009, Storey 2015). Whether urchins can transition to hypometabolic states in low-productivity barrens and how this effect might scale with body size and/or biomass remains, to our knowledge, untested.

We hypothesized that emaciated *M. franciscanus* individuals in barrens dramatically depress their metabolism to maximize energetic efficiency for survival. To quantify the metabolic state of *M. franciscanus* without the confounding influences of locomotive activity and postprandial effects, we targeted the resting metabolic rate (RMR). Specifically, we hypothesized that the body size-specific RMR (i.e. RMR for a given body mass or body volume) for *M. franciscanus* in food-depauperate barrens would be lower relative to kelp forest habitats. To test for an effect of habitat on RMR, we compared individuals spanning small (minimum: 25 mm test diameter) through large (maximum: 138 mm test diameter) body sizes living in kelp forests and barrens in BC, Canada (Fig. 1) in May 2019 on Quadra Island and July 2019 in Haida Gwaii. Study sites included rocky subtidal kelp forests (approx. 2 m below mean low water) and barrens (approx. 12 m below mean low water) at three locations, Faraday (52.61°N, 131.49°W) and Murchison (52.60°N, 131.45°W) in Gwaii Haanas on Haida Gwaii, and Surge Narrows (50.22°N, 125.16 °W) between Quadra and Maurelle Islands. Based on field observations, we expected size-specific reductions in gonadal mass associated with food limitation in barrens (Fig. 1) giving rise to different allometric exponents for the relationship between body size and gonadal mass in kelp forests relative to barrens (Ebert et al. 2011).

**Figure 1.**
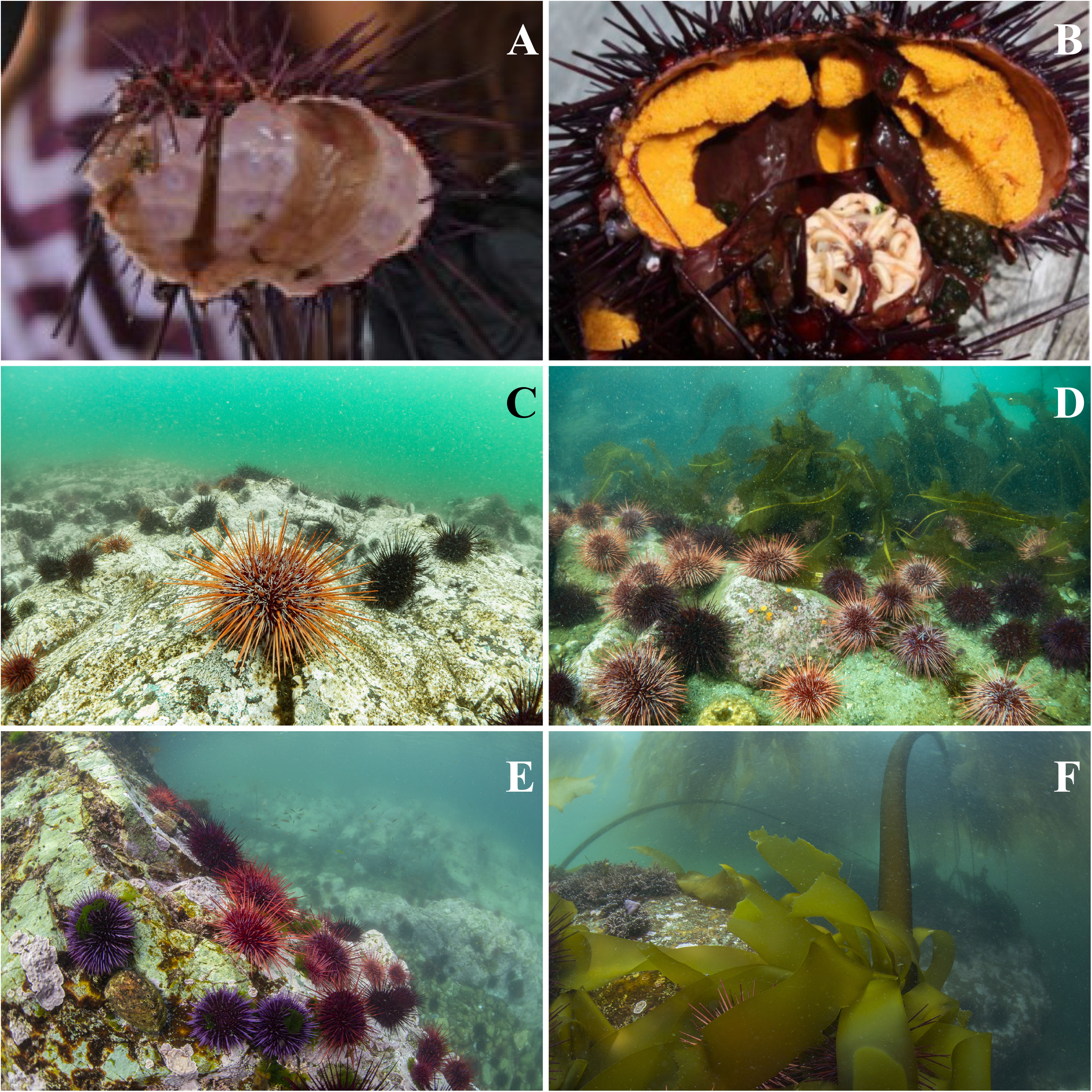
Visual comparison of typical qualitative habitat characteristics and internal anatomy of resident sea urchins in kelp forest versus barrens habitats. A) Dissected urchins with diminutive gonads typical of barrens. B) Dissected urchins with robust gonads typical of kelp forest urchins. C) Barrens habitat at Surge Narrows, BC, at a depth of approx. 12 m below chart datum low tide. D) At the edge of kelp forest habitat at Surge Narrows, BC, Canada, at a depth of approx. 3 m relative to chart datum low tide. E) Barrens habitat at Murchison Island in Gwaii Haanas at a depth of approx. 12 m relative to chart datum low tide. F) At the edge of kelp forest habitat at Murchison Island in Gwaii Haanas National Park Reserve, National Marine Conservation Area Reserve, and Haida Heritage Site, at a depth of approx. 3 m relative to chart datum low tide. Photo A was taken by Spindel, B was taken by Lee, C and D were taken by Markus Thompson, and E and F were taken by Ryan Miller.

We measured respiration rates of *M. franciscanus* in a cumulative 70 individuals from kelp forest habitats and 79 individuals from barrens across our three sites as a proxy for metabolic rate using custom-built sealed chambers (Appendix Fig. S1) fitted with flow-through optical oxygen sensors and a temperature sensor (Presens Precision Sensing GmbH). To quantify RMR, we measured respiration rates after a 48-hour period of starvation post-collection from the wild following (Lighton 2018). We conducted quality control on oxygen time series data using the R package respR (Harianto et al. 2019). To contextualize metabolic rates, we measured body size (i.e. internal urchin test volume), total biomass, and gonadal mass. We calculated internal urchin test volume (V) from test height (H) and test diameter (D) assuming oblate spheroid geometry (V = 4/3π D^2^H) and recorded two metrics of whole urchin biomass: first wet mass then ash-free dry mass (AFDM). To measure AFDM, we first dried samples for 24 hours at 60□ in a drying oven (Thermo Scientific) then combusted dry samples for six hours at 450□ in a muffle furnace (Thermo Scientific). AFDM targets non-skeletal soft tissue quantified as the difference between dry mass and post-combustion ash mass. We measured whole-animal wet mass after cracking and discarding seawater from inside the test, and gonadal wet mass following 30 seconds of drying dissected gonads on a paper towel to correct for variation in water content. Only urchins from Surge Narrows had AFDM measured because of equipment availability. We estimated additive and interactive effects of habitat and site and body size/mass on RMR and gonadal wet mass by fitting the metabolic scaling function (i.e. *RMR — aB^β^* — exp [log(α) + /βlog (β)]) using generalized linear mixed effects models fitted using the R package glmmTMB (Brooks et al. 2017) with a lognormal likelihood, treating habitat, log-scale metrics of body size/mass and site as fixed effects, and date and respiration chamber as random effects (Fig. 2).

**Figure 2.**
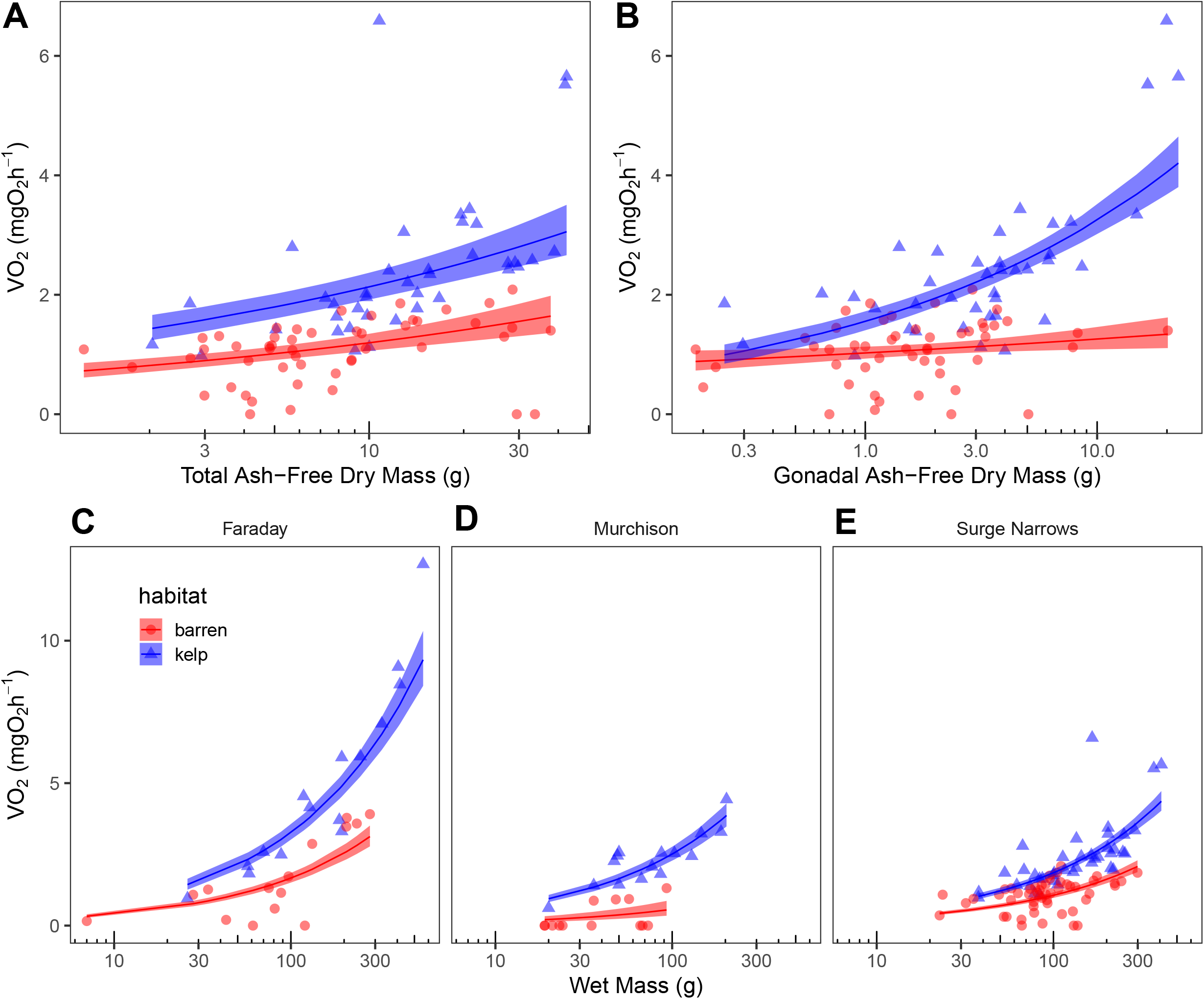
Comparison of resting metabolic rate in *M. franciscanus* versus metrics of body size by habitat and site. Dots represent 2019 volumetric oxygen consumption rate (VO_2_) measurements from individual urchins. Lines and ribbons represent modelled mean VO_2_ and SE, respectively. Panels A and B contrast the scaling relationship in barren versus kelp forest habitat at Surge Narrows, BC, Canada, between VO2 and total ash-free dry mass and gonadal ash-free dry mass, respectively. Panels C-E show geographic comparison of the scaling relationships in barrens versus kelp forest habitats between VO_2_ and body size (i.e. test volume) among sites at Faraday Island, Murchison Island, and Surge Narrows, BC, Canada.

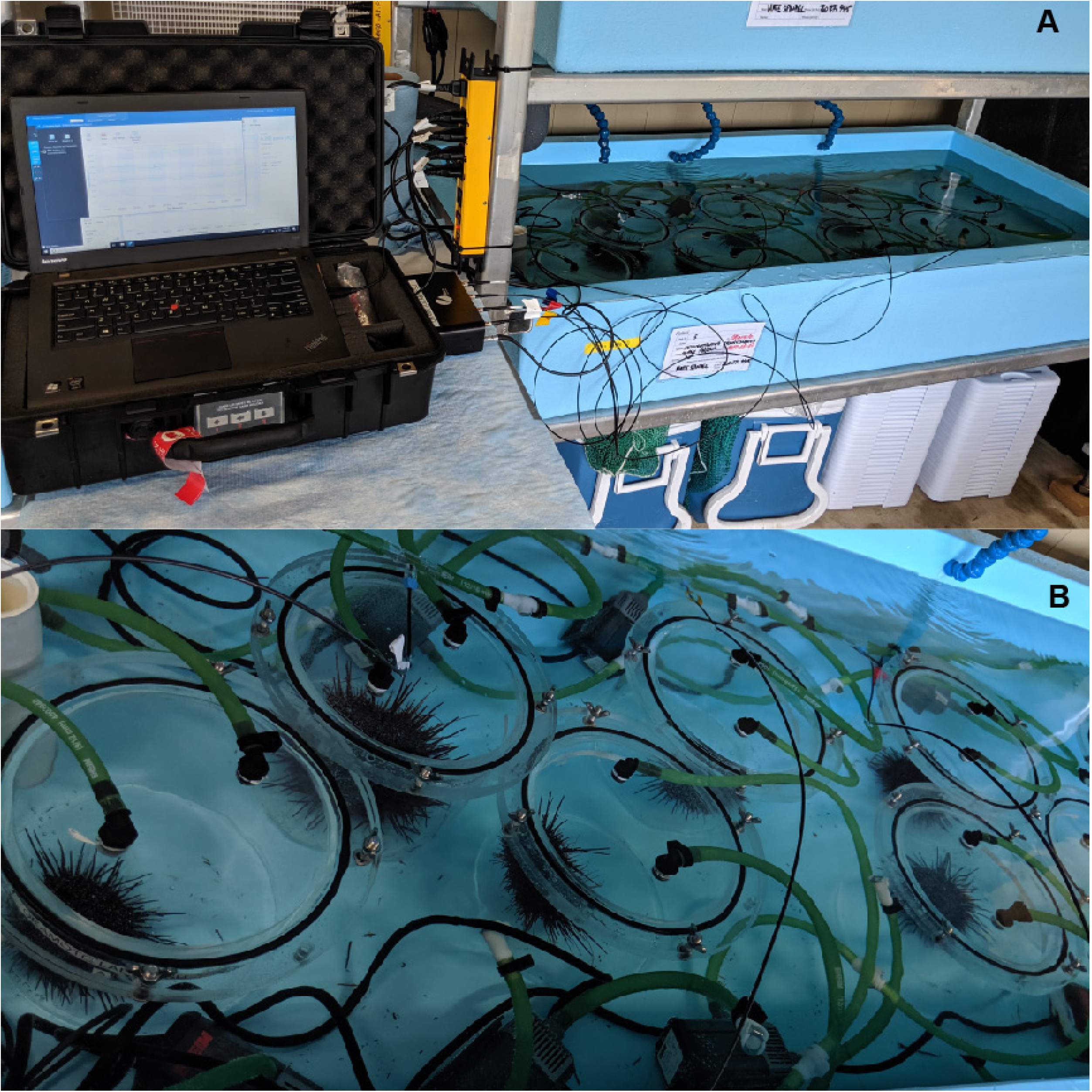

*M. franciscanus* in barrens exhibited dramatically depressed metabolic rates compared to animals in kelp forests, even after accounting for wet body mass, body volume, or AFDM (Fig. 2). For a given whole-animal wet mass, RMR was nearly 50% lower in barrens urchins (multiplicative effect on the natural scale was 0.51, 95% CI: 0.43-0.61, compared to kelp forests, χ^2^_df=1_ = 59.06, P < 0.001). When scaled to wet mass, RMR varied by site (χ^2^_df=2_ = 14.38, P = 0.001), but we found no significant interactions between habitat, wet mass, or site (Appendix S1: Table S6). For a given test volume, RMR was 56% lower in barrens urchins (multiplicative effect on the natural scale was 0.44, 95% CI: 0.38-0.52, compared to kelp forests, χ^2^_df=1_ = 104.48, P < 0.001). When scaled to body volume, RMR varied by site (χ^2df=2^ = 18.63, P < 0.001), but we found no significant interactions among site, habitat, or body volume (Appendix S1: Table S8). Urchins at all three sites had significantly lower gonadal wet mass in barrens relative to kelp forest habitats as indicated by the test-volume x habitat interaction (χ^2^_df=1_= 26.48, P < 0.001) with, on average 44.6% percent lower gonad mass in barrens. At Surge Narrows (where lab facilities allowed us to take metrics of AFDM) RMR was 43% lower in barrens versus kelp forest habitats when scaled to metabolically active body mass (multiplicative effect on the natural scale was 0.57, 95% CI: 0.38-0.86, compared to kelp forests, χ^2^_df=1_ = 7.35, P = 0.01). For a given gonadal AFDM, RMR was 34.0% lower in barrens urchins (multiplicative effect on the natural scale was 0.66, 95% CI: 0.53-0.82, compared to kelp forests, χ^2^_df=1_ = 14.18, P < 0.001). There was no significant interaction between total AFDM and habitat (RMR versus log total AFDM x habitat: χ^2^_df=1_ = 0.01, P = 0.92) but there was a significant interaction between gonadal AFDM and habitat (RMR versus log gonadal AFDM x habitat: χ^2^_df=1_ = 7.08, P = 0.01).

These observations demonstrate that *M. franciscanus* in barrens not only have reduced gonadal reserves and thus lower metabolically active body mass, but also exhibit substantial reductions in mass-specific RMR. Whole-animal RMR was higher for larger individuals in both habitats at Faraday Island relative to the other two sites, but barren urchins were still metabolically depressed relative to adjacent kelp forest urchins. This site-specific difference was likely due to greater food availability, potentially as a result of increased algal subsidy from exogenous sources at Faraday relative to the other two sites. Large urchins in both habitats at Faraday had higher gonadal mass, indicating higher food availability (Rogers-Bennett and Okamoto 2020) than large urchins in the other two sites despite barrens having no endogenous, macroscopic algae beyond encrusting corallines (Spindel, Okamoto, Lee, unpublished data). Reductions in metabolic rate substantially exceeded that expected by changes in body mass alone. One plausible explanation for reductions in mass-specific RMR in *M. franciscanus* is a reduction in cellular metabolism. Based on evidence from mammalian and avian species, one might expect the nature of this hypometabolism would depend on the frequency and/or duration of food deprivation in an organism’s past (McCue et al. 2017). For example, starvation may “reprogram” fetal humans in utero via epigenetic effects so they develop metabolic syndrome in adulthood (Rinaudo and Wang 2012). Another plausible explanation is that the proportion of tissues with lower metabolic rate increases by utilizing reversible biomass. Evidence from mammalian (Rolfe and Brown 1997) and avian (Daan et al. 1990) species shows that metabolic rates differ among tissue types. For example, liver and gastrointestinal tissue contribute equivalent body mass percentages in both humans and rats, but metabolic contributions of these tissues differ widely (17% versus 10% in humans, and 20% versus 5% in rats, respectively) (Rolfe and Brown 1997). Therefore, changes in body composition alone can theoretically produce changes in whole-animal metabolic rates. As they deplete lipid-rich reserves that may have lower maintenance costs than other tissues, animals must either reduce their locomotive activity and/or depress cellular metabolism to endure the energetic burden of food deprivation. The predominant source of change in body composition we observed in metabolically depressed urchins was a reduction in gonad mass. One would expect gonadal tissue would be less metabolically active relative to other tissue types, although we did not measure tissue-specific respiration rates. A reduction in less-metabolically active tissues would be unlikely to explain whole animal biomass-specific reductions in RMR. Therefore, we submit that a greater proportion of the observed metabolic depression is likely due to regulation at a cellular level rather than shifts in tissue composition, but further studies are required to assess this hypothesis.

This phenomenon of mass-specific metabolic depression may help individuals balance growth and survival amid collapses in or intermittent availability of food. Despite these changes, urchins can capitalize on any newly available food in short order. Laboratory studies showed that starved purple urchins (Okamoto 2014), and red urchins (Spindel & Okamoto, unpublished data) can recover their reduced gonad mass two to three months after re-feeding and revert in a similar time frame. Unlike many species that cope with food deprivation by entering a dormancy phase including metabolic depression and suspended development (Hand and Hardewig 1996, Hand et al. 2016), *M. franciscanus* continues to grow (albeit more slowly), move, and opportunistically feed while metabolically depressed (Okamoto, Spindel, Lee, unpublished tag-recapture data).

Controlled experiments are required to characterize how and why this may occur, over what time scales, and to evaluate impacts of metabolic depression on rates of herbivory and the persistence of barrens. However, these observations from three sites in BC support the notion that *M. franciscanus* in barrens can dramatically reduce their energetic demands. Moreover, these shifts in metabolic rate may provide a mechanism that facilitates barren state stability over long time scales as *M. franciscanus* can lower energetic demands while they wait for small pulses of food, scrape by on low-productivity resources, and suppress recruitment of macroalgae for months, years, or decades.

## ACKNOWLEDGEMENTS

We thank Leandre Vigneault of Marine Toad Enterprises Inc. and Dan McNeill, Ben Penna, Jaasaljuus Yakgujanaas, Vanessa Bellis and Dion Lewis of the Council of the Haida Nation Haida Fisheries Program for assistance with specimen collection in Gwaii Haanas; Markus Thompson of Thalassia Environmental, Ryan Miller of Miller Marine Diving Services, and Ondine Pontier of the Hakai Institute for assistance with specimen collection and underwater photography at Quadra Island; Eric Peterson and the Hakai Institute for providing laboratory facilities at Quadra Island and for technical support; and Gwaii Haanas National Park Reserve, National Marine Conservation Area Reserve, and Haida Heritage Site for technical support, remote infrastructure and field personnel including Charlotte Houston, Niisii Guujaaw, Clint Johnson Kendrick, Chavonne Guthrie and Marilyn Deschenes. This study was funded by an FSU APACT grant to DKO, Parks Canada Conservation and Restoration (CoRe) project funding to the Gwaii Haanas Field Unit with LL as technical lead and DKO via contribution agreement with FSU, the William R. and Lenore Mote Eminent Scholar in Marine Biology Endowment at FSU to NBS, a Professional Association of Diving Instructors (PADI) Foundation Research Grant to NBS, an Academy of Underwater Arts and Sciences Zale Parry Scholarship to NBS, and the Tula Foundation.

