## Appendix for "Metabolic depression in sea urchin barrens associated with food deprivation"

APPENDIX S1

Table S1: GLMM summary for $VO_2$ regressed against total AFDM by habitat at Surge Narrows.

| | $VO_2$ | | |
|---|---|---|---|
| *Parameter* | *Estimates* | *95% CI* | *p* |
| **(Intercept)** | **-0.37** | **-0.72 – -0.02** | **0.040** |
| **whole.afdm.g.approx [log]** | **0.24** | **0.08 – 0.40** | **0.003** |
| **habitat [kelp]** | **0.56** | **0.15 – 0.96** | **0.007** |
| whole.afdm.g.approx [log] * habitat [kelp] | 0.01 | -0.18 – 0.20 | 0.919 |
| **Random Effects** | | | |
| $\sigma^2$ | | | |
| $\tau_{00}$ chamber_id:date | 0.08 | | |
| $\tau_{00}$ date | 0.01 | | |
| N chamber_id | 6 | | |
| N date | 3 | | |
| Observations | 89 | | |

Table S2: Analysis of deviance (type III Wald $\chi^2$ tests) results for the additive and interactive effects of total AFDM and habitat on $VO_2$ at Surge Narrows.

| | $\chi^2$ | Df | P |
|---|---|---|---|
| **log(whole.afdm.g.approx)** | **8.596** | **1** | **0.003** |
| **habitat** | **7.354** | **1** | **0.007** |
| log(whole.afdm.g.approx):habitat | 0.010 | 1 | 0.919 |

Table S3: GLMM summary for $VO_2$ regressed against gonadal AFDM by habitat at Surge Narrows.

| | | **$VO_2$** | |
|---|---|---|---|
| *Parameter* | *Estimates* | *95% CI* | *p* |
| (Intercept) | 0.03 | -0.16 – 0.22 | 0.792 |
| gonad.afdm.g.total.approx [log] | 0.09 | -0.05 – 0.23 | 0.218 |
| **habitat [kelp]** | **0.42** | **0.20 – 0.63** | **<0.001** |
| **gonad.afdm.g.total.approx[log] * habitat [kelp]** | **0.23** | **0.06 – 0.40** | **0.008** |
| **Random Effects** | | | |
| $\sigma^2$ | 0.23 | | |
| $\tau_{00\ chamber\_id:date}$ | 0.04 | | |
| $\tau_{00\ date}$ | 0.00 | | |
| ICC | 0.14 | | |
| $N_{chamber\_id}$ | 6 | | |
| $N_{date}$ | 3 | | |
| Observations | 89 | | |
| Marginal $R^2$ / Conditional $R^2$ | 0.404 / 0.490 | | |

Table S4: Analysis of deviance (type III Wald $\chi^2$ tests) results for the additive and interactive effects of gonadal AFDM and habitat on $VO_2$ at Surge Narrows.

| | $\chi^2$ | Df | P |
|---|---|---|---|
| log(gonad.afdm.g.total.approx) | 1.516 | 1 | 0.218 |
| **habitat** | **14.179** | **1** | **0.000** |
| **log(gonad.afdm.g.total.approx):habitat** | **7.078** | **1** | **0.008** |

Table S5: GLMM summary for $VO_2$ regressed against whole-animal wet mass by habitat and site.

| | $VO_2$ | | |
|---|---|---|---|
| *Parameter* | *Estimates* | *95% CI* | *p* |
| **(Intercept)** | **-2.29** | **-2.85 – -1.74** | **<0.001** |
| **animal_wet_mass_g [log]** | **0.61** | **0.50 – 0.72** | **<0.001** |
| **habitat [kelp]** | **0.67** | **0.50 – 0.84** | **<0.001** |
| **site [Murchison]** | **-1.07** | **-1.99 – -0.15** | **0.023** |
| **site [Surge Narrows]** | **-0.45** | **-0.71 – -0.18** | **0.001** |
| habitat [kelp] * site[Murchison] | 0.81 | -0.10 – 1.71 | 0.081 |
| habitat [kelp] * site[Surge Narrows] | -0.12 | -0.36 – 0.13 | 0.348 |
| **Random Effects** | | | |
| $\sigma^2$ | 0.35 | | |
| $\tau_{00\ chamber\_id:date}$ | 0.02 | | |
| $\tau_{00\ date}$ | 0.00 | | |
| $N_{chamber\_id}$ | 8 | | |
| $N_{date}$ | 5 | | |
| Observations | 149 | | |
| Marginal $R^2$ / Conditional $R^2$ | 0.620 / NA | | |

Table S6: Analysis of deviance (type III Wald $\chi^2$ tests) results for the additive and interactive effects of total wet mass, habitat, and site on $VO_2$.

| | $\chi^2$ | Df | P |
|---|---|---|---|
| **log(animal_wet_mass_g)** | **112.757** | **1** | **<0.001** |
| **habitat** | **59.056** | **1** | **<0.001** |
| **site** | **14.378** | **2** | **0.001** |
| habitat:site | 4.32 | 2 | 0.115 |

Table S7: GLMM summary for $VO_2$ regressed against test volume by habitat and site.

| | **$VO_2$** | | |
|---|---|---|---|
| *Parameter* | *Estimates* | *95% CI* | *p* |
| **(Intercept)** | **-1.99** | **-2.42 – -1.56** | **<0.001** |
| **Spheroid.volume.ml [log]** | **0.53** | **0.45 – 0.61** | **<0.001** |
| **habitat [kelp]** | **0.82** | **0.66 – 0.97** | **<0.001** |
| **site [Murchison]** | **-0.97** | **-1.81 - -0.12** | **0.024** |
| **site [Surge Narrows]** | **-0.49** | **-0.73 - -0.25** | **<0.001** |
| habitat [kelp] * site[Murchison] | 0.57 | -0.27 – 1.40 | 0.186 |
| habitat [kelp] * site[Surge Narrows] | -0.20 | -0.43 – 0.04 | 0.099 |
| **Random Effects** | | | |
| $\sigma^2$ | 0.33 | | |
| $\tau_{00}$ chamber_id:date | 0.01 | | |
| $\tau_{00}$ date | 0.00 | | |
| N chamber_id | 8 | | |
| N date | 5 | | |
| Observations | 149 | | |
| Marginal $R^2$ / Conditional $R^2$ | 0.653 / NA | | |

Table S8: Analysis of deviance (type III Wald $\chi^2$ tests) results for the additive and interactive effects of test volume, habitat, and site on $VO_2$.

| | $\chi^2$ | Df | P |
|---|---|---|---|
| **log(spheroid.volume.ml)** | **160.199** | **1** | **<0.001** |
| **habitat** | **104.484** | **1** | **<0.001** |
| **site** | **18.632** | **2** | **<0.001** |
| habitat:site | 5.015 | 2 | 0.082 |

Table S9: GLMM summary for gonadal wet mass regressed against test volume by habitat and site.

| | Gonadal Wet Mass | | |
|---|---|---|---|
| *ParameterParameter* | *Estimates* | *95% CI* | *p* |
| **(Intercept)** | **-0.97** | **-1.87 – -0.07** | **0.035** |
| **spheroid.volume.ml [log]** | **0.63** | **0.46 – 0.80** | **<0.001** |
| **habitat [kelp]** | **-4.1** | **-5.92 – -2.28** | **<0.001** |
| **site [Murchison]** | **0.39** | **0.04 – 0.74** | **0.028** |
| site [Surge Narrows] | 0.2 | -0.01 – 0.41 | 0.061 |
| **spheroid.volume.ml [log] * habitat [kelp]** | **0.81** | **0.50 – 1.12** | **<0.001** |
| Observations | 149 | | |
| $R^2$ conditional / $R^2$ marginal | NA / 0.470 | | |

Table S10: Analysis of deviance (type III Wald $\chi^2$ tests) results for the additive and interactive effects of test volume, habitat, and site on gonadal wet mass.

| | $\chi^2$ | Df | P |
|---|---|---|---|
| **log(spheroid.volume.ml)** | **54.213** | **1** | **0** |
| **habitat** | **19.487** | **1** | **0** |
| site | 5.399 | 2 | 0.0672 |
| **log(spheroid.volume.ml):habitat** | **26.478** | **1** | **0** |

### FIGURE CAPTIONS

Figure S1. Photographs of respirometry apparatus. Chambers are custom-built five-liter acrylic cylinders sealed with o rings. We measured oxygen concentration using optical sensors affixed to flow-through cells and temperature using a temperature probe (Presens Precision Sensing GmbH). We logged oxygen concentration and temperature time series using a four-channel respirometer paired with a PC laptop running PreSens Measurement Studio 2 software. A) Complete respirometry array including two sets of four respiration chambers, one set for acclimation, one for measurement of respiration. B) Close-up of respiration chambers containing *M. franciscanus*.
